# Biogeographic position and body size jointly set lower thermal limits of wandering spiders

**DOI:** 10.1101/2021.01.04.422639

**Authors:** Jérémy Monsimet, Hervé Colinet, Olivier Devineau, Denis Lafage, Julien Pétillon

**Affiliations:** Department of Forestry and Wildlife management, Inland Norway University of Applied Sciences, Campus Evenstad, Koppang, Norway; UMR CNRS 6553 ECOBIO, Université de Rennes, Rennes, France; Department of Environmental and Life Sciences/Biology, Karlstad University, Karlstad, Sweden

**Author notes:** Correspondence: Jérémy Monsimet < >.

**Keywords:** Supercooling ability, fishing spiders, freezing, climate change, *Dolomedes*

## Abstract

Most species encounter large variations in abiotic conditions along their distribution range. Climate, and in particular temperature, varies along clinal gradients, which determines phenotypic plasticity, local adaptations and associated physiological responses of most terrestrial ectotherms, such as insects and spiders. This study aimed to determine how the biogeographic position of populations and the body size of two wandering spiders set their limits of cold (freezing) resistance. Using an ad-hoc design, we sampled relatively large numbers of individuals from four populations of *Dolomedes fimbriatus* and one population of the sister species *Dolomedes plantarius* originating from contrasting climatic areas (temperate and continental climate), and compared their supercooling ability as an indicator of cold resistance. Results indicated that spiders from northern (continental) populations had higher cold resistance than spiders from a southern (temperate) populations. Larger spiders had a lower supercooling ability in northern populations. The red-listed and rarest *D. plantarius* was slightly less cold-tolerant than the more common *D. fimbriatus*, and this might be of importance in a context of climate change that could imply colder overwintering habitats in the north due to reduced snow cover protection.

## Introduction

The ability of a species to cope with variations in abiotic conditions influences its distribution range (Gaston 2003). Abiotic factors, and among them temperature, shape the geographic range of ectotherm species, and this is even more relevant in the context of global warming (Somero 2012, Addo-Bediako et al. 2000). Some ecthoterms survive freezing and are freeze tolerant whereas other ectotherms are freeze intolerant. Freezing tolerant species, like some alpine species, tend to freeze at relatively high subzero temperatures with ice nucleators and cryoprotectants, inducing and protecting against freezing stress respectively, instead of having high supercooling abilities, i.e. low supercooling point (SCP) (Duman 2001, Bale 2002, Duman et al. 2004). Freeze intolerant arthropods, which include freeze-avoidant, chill tolerant, chill-susceptible and opportunistic-survival classes, can exhibit deep supercooling ability, ranging from −15 to −25°C (Danks 2004), by producing polyols and antifreeze proteins (Duman 2001, Bale 2002).

Many different measures are used to illustrate the thermal performance of populations (Sinclair et al. 2015). It could be depicted by a thermal performance curve representing how a temperature gradient influences arthropod activity (Sinclair et al. 2012, 2015). As the estimation of thermal performances is influenced by many factors such as phenotypic plasticity (Schulte et al. 2011) or evolutionary adaptation (Jensen et al. 2019), measuring an anchor point like the SCP is useful to assess the cold tolerance class of species. Indeed, the SCP represents the lower lethal temperature (LLT) for freezing-avoidant species and is still a useful indicator for chill-tolerant species as SCP and LLT are almost similar for them (Bale 1996). However, many ectotherms classified as chill-susceptible or opportunistic-survival, die at temperatures well above SCP, the latter being less resistant than the former (Overgaard and MacMillan 2017, Bale 2002). Even though the ecological value of the SCP has been debated (Renault et al. 2002, Ditrich et al. 2018), it is still a useful metric to explore and describe the cold tolerance strategy of poorly studied species, such as spiders (Sinclair et al. 2015).

Latitude and winter conditions influence the temperature gap between the SCP and the lower lethal temperature (Addo-Bediako et al. 2000, Vernon and Vannier 2002). Indeed, based on cold hardiness strategies defined by Bale (1996), opportunistic-survival animals are mainly found in tropical and semi-tropical regions, chill susceptible and chill-tolerant in temperate and sub-polar regions and freeze avoidant in region with severe cold winter conditions.

Body size influences and is influenced by the animal’s stage, its body fat content or the concentration of ice-nucleating bacteria, which affect the SCP (David and Vannier 1996, Johnston and Lee 1990, Colinet et al. 2007). The size of animals also changes along latitudinal and altitudinal clines. Both an increase and a decrease of body size towards northern latitude were observed and theorised under the Bergmann and converse Bergmann rules respectively (Blanckenhorn and Demont 2004). For ectotherms, these two rules were first opposed (Voorhies 1996, Mousseau 1997) but it seems that both larger and smaller individuals at northern latitudes is possible and the two rules are eventually not exclusive (Blanckenhorn and Demont 2004), possibly co-existing in close species (e.g. in artic wolf spiders, see Ameline et al. 2018). The latitudinal size cline is of importance as body size also influences cold hardiness (Ansart et al. 2014), e.g. with smaller arthropods having better supercooling capabilities than larger ones (Sømme 1982, David et al. 1996, Colinet et al. 2007, Sinclair et al. 2009). Hence, a negative relationship between ectotherms size and the ability to supercool has been reported (Lee and Costanzo 1998). Consequently, smaller individuals could benefit from colder temperatures under harsher winter conditions at northern latitudes.

Most studies investigating latitudinal clinal changes of arthropods’ physiological tolerance focused on differences between species rather than among populations of the same species (Spicer and Gaston 1999 but see, e.g. Jensen et al. 2019). Physiological tolerance is a basal trait in arthropods, but it has evolved many times (Sinclair et al. 2003). Most of the knowledge on cold tolerance of arthropods comes from the study of insects, and different mechanisms might influence the cold hardiness of insects and arachnids. Indeed, Anthony and Sinclair (2019) showed divergent cryoprotective dehydration, the action of losing water by evaporation at low temperature, between insects and arachnids and the absence of coma under hypoxic conditions is also remarkable in spiders (Pétillon et al. 2009). To our knowledge, not all spiders are freezing tolerant (Nentwig 2012). The same cold hardiness classes are used to categorise freezing intolerance of spiders and insects. Indeed, some spiders are freeze-avoidant, others chill-tolerant or chill susceptible (Kirchner 1973, Anthony et al. 2019).

Although latitudinal variations in the cold hardiness of arachnids have been the subject of recent attention (e.g. Anthony et al. 2019), studies comparing populations within species are lacking (but see e.g. Murphy et al. 2008), this despite the recognised importance of comparative approaches (e.g. see Ansart et al. 2014). Tough sampling conditions at high latitude in the northern hemisphere may limit sampling of a sufficient number of individuals and thus prevent studies from considering the northern part of a species range.

In this study, we assessed the variation in cold resistance, estimated through SCP ability of different populations and species of fishing spiders (Araneae, Pisauridae) with contrasted distributions. We hypothesised that (i) northern populations of *Dolomedes fimbriatus* have lower SCP values than southern populations, (ii) the size of spider in the north is positively related to the SCP, and (iii) the species reaching the northern latitudes (here *D. fimbriatus*) has lower SCP values than the more southern limited species (*D. plantarius*: Monsimet et al. 2020), potentially due to their relatively smaller body size. These hypotheses were experimentally tested in two European *Dolomedes* species using relatively high numbers of field-collected spiders for representative results and robust statistics.

## Materials and Methods

### Case study species and sampling locations

The fishing spiders, *Dolomedes plantarius* and *Dolomedes fimbriatus* are widespread in Europe with a northern range limit in Fennoscandia. *D. plantarius* has a lower population density and is red-listed at the European scale (World Conservation Monitoring Centre 1996). The latitudinal contrast encompassed two different biogeographic positions, characterizing two different climatic areas (continental, coded C hereafter *versus* temperate, coded T). Individuals sampled at their range limit were compared with others from a central latitude of the distribution. We sampled two sites with *D. fimbriatus* and one site with *D. plantarius* in Fennoscandia (C1, C2 and C3; fig. 1), which characterise the northern population, subject to a continental climate. In addition, we sampled two sites with *D. fimbriatus* in France (T1 and T2; fig. 1), representing the centrally distributed populations exposed to a temperate climate. Given the conservation status of *D. plantarius* in Europe, we chose to limit our sampling of this species to the area where it is most abundant (Fennoscandia).

**Figure 1:**
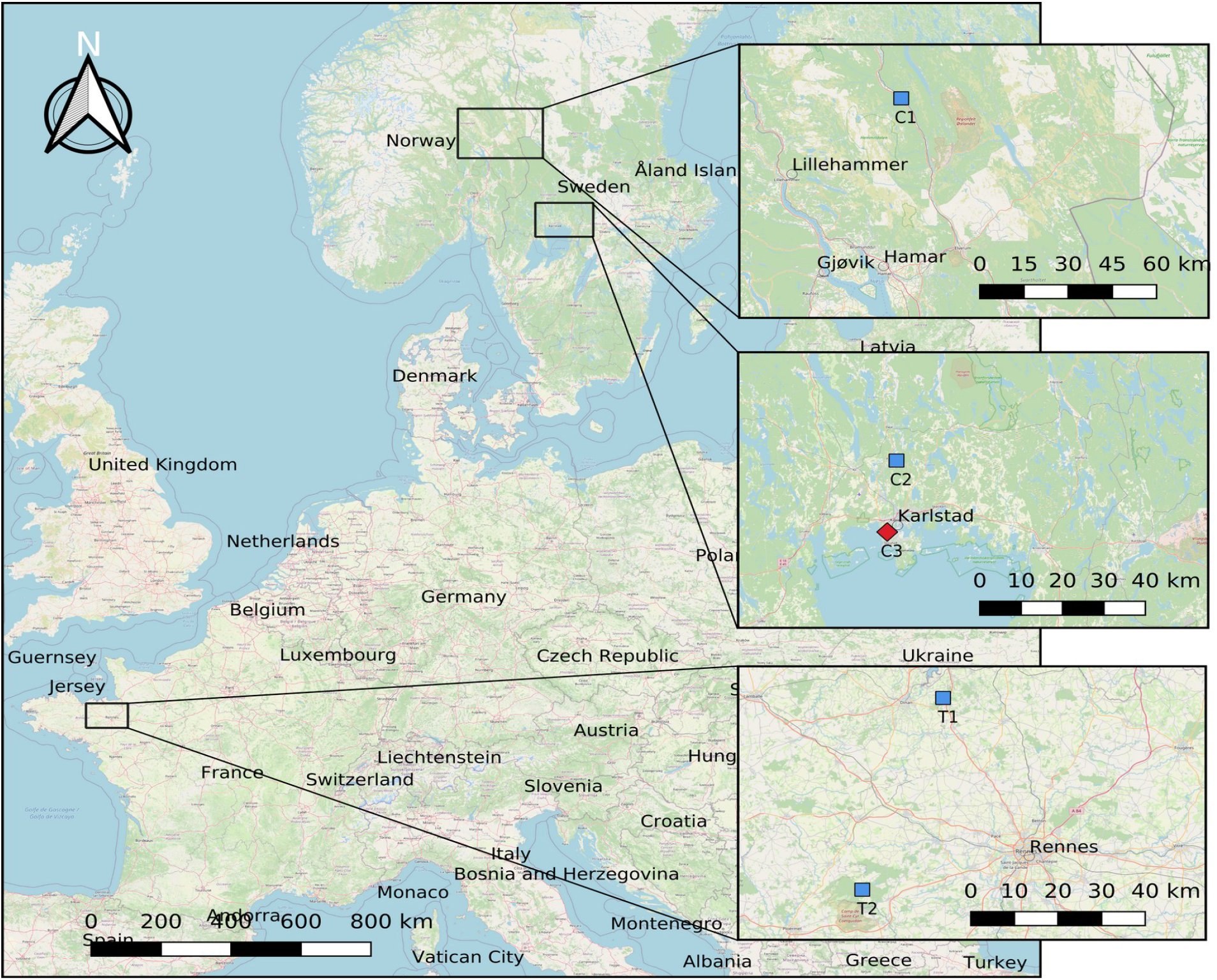
Location of sampling sites for *Dolomedes fimbriatus* (blue squares) and *Dolomedes plantarius* (red square) in France and Fennoscandia.

As the SCP is influenced by the developmental stage (Aitchison 1984, Anthony et al. 2019), we sampled only juvenile spiders of both sexes. The peak of the breeding season of European *Dolomedes* is in late July (Smith 2000). Females keep egg sacs several weeks before building a nursery web where eggs will hatch and from which spiderlings will later spread out into the surroundings. Juvenile spiders overwinter, but not adults, similarly to other species in the genus (Guarisco 2010). We sampled *D. fimbriatus* by sweep-netting the vegetation on sunny and windless days. We sampled *D. plantarius* on the water surface by visual hunting, and active hunting by perturbing the water surface. We sampled, and latter tested the SCP of about 24 spiders at each sampling site (*n* = 24,24,21,26,24 for C1,C2,C3,T1,T2 respectively, table 1).

**Table 1:** Description of the climatic conditions at the sampling sites, based on the Köppen-Geiger climate classification (Kottek et al. 2006). N: number of spiders tested; SCP: mean SCP ± SD; Length: mean length of the carapace +-SD; Mean temp: annual mean temperature; Diurnal range: mean diurnal range (extracted from Fick and Hijmans (2017)).

| Sites | Species | N | Country | Climate | SCP (°C) | Length (mm) | Mean Temp | Diurnal range |
| --- | --- | --- | --- | --- | --- | --- | --- | --- |
| C1 | *D. fimbriatus* | 24 | Norway | Continental | -9.08±0.45 | 4.13±0.52 | 2.56 | 9.50 |
| C2 | *D. fimbriatus* | 24 | Sweden | Continental | -9.06±0.4 | 4.43±0.56 | 5.52 | 8.54 |
| C3 | *D. plantarius* | 21 | Sweden | Continental | -7.56±0.32 | 5.36±0.69 | 6.05 | 7.78 |
| T1 | *D. fimbriatus* | 26 | France | Temperate | -7.78±0.4 | 4.62±0.46 | 11.62 | 7.03 |
| T2 | *D. fimbriatus* | 24 | France | Temperate | -5.39±0.4 | 4.44±0.48 | 11.14 | 6.30 |

### Measurement of the supercooling point

To determine the SCP, we placed the spiders in centrifuge tubes, which were submerged in a cryostat bath (Polystat CC3, Huber Kältemaschinenbau AG, Germany) filled with heat transfer fluid (Thermofluid SilOil, Huber, Germany). The temperature of the bath was slowly reduced at a rate of 0.5°C min^−1^ to reach a target temperature of − 30°C. To monitor the temperature of the spiders, we placed a K-type thermocouple in direct contact with the spider opisthosoma, secured with Parafilm® and connected to a Testo 175T3 temperature data logger (Testo SE& Co., Germany). We recorded the temperature every ten seconds. The SCP was defined as the temperature at the onset of the freezing exotherm produced by the latent heat (see fig. 2 for representative exotherms).

**Figure 2:**
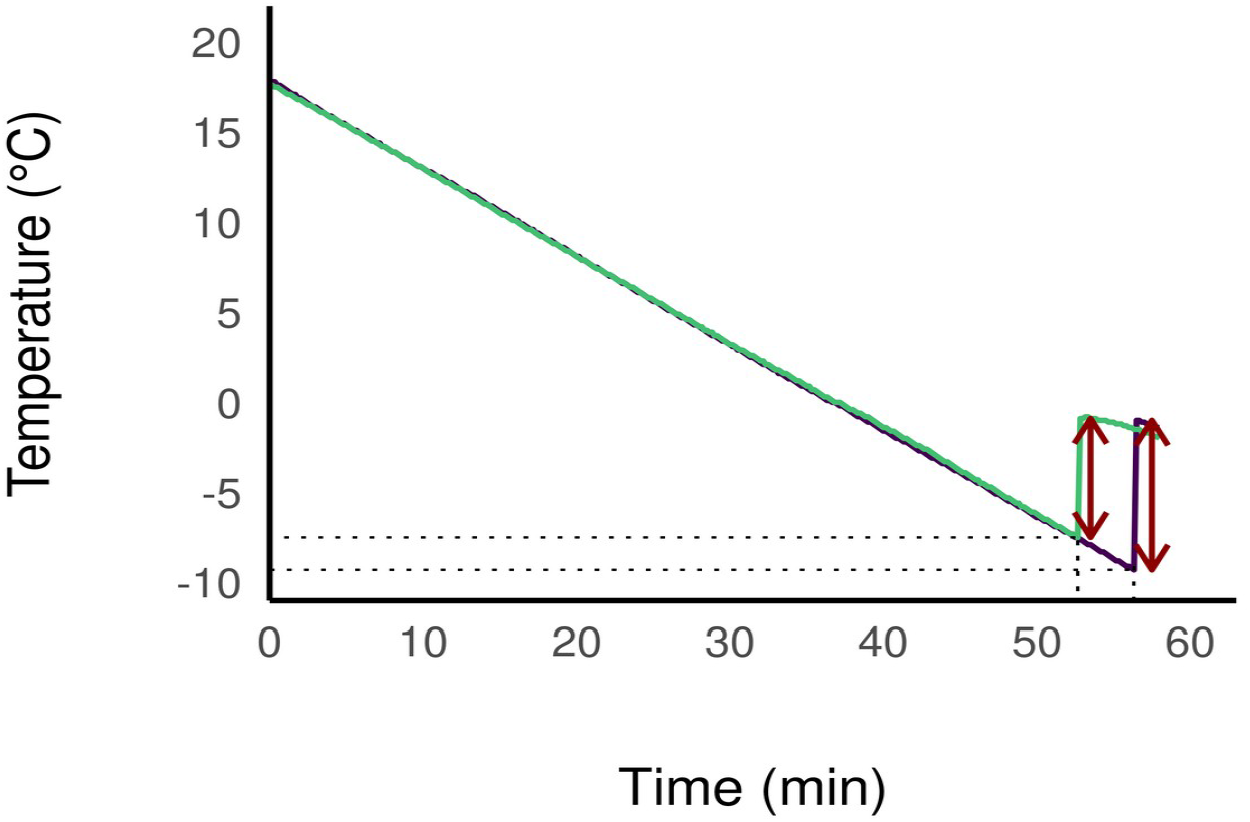
Cooling curves of *D. plantarius* (one spider from C3, in yellow) and *D. fimbriatus* (one spider from C2, in purple) recorded during a cooling experiment. The SCP (dotted line) is followed by the exotherm (dark-red arrows), a sudden increase in the measured temperature due to the release of latent heat linked to the phase change during freezing.

As the number of spiders tested per day was limited by the capacity of the instrument (4 spiders at a time), we later accounted for the time lag between capture and test in our models (variable Diff).

### Measurement of spider body size

We measured the spiders’ body size after the SCP experiment to avoid injuring the spiders and biasing the results. We took a picture of the spider’ back together with a measuring tape for measuring the body size later in the ImageJ software (Schneider et al. 2012). We measured the highest length and largest width of the carapace (prosoma) which are commonly used as proxy for whole body size, fitness and metabolic rate in spiders (Jakob et al. 1996, Penell et al. 2018).

### Data treatment

The carapace width and length were highly correlated (rho = 0.83, Pearson correlation test), so we used the carapace length as a proxy of body size (Jakob et al. 1996) and referred to as body size hereafter.

### Comparison of SCP across latitudes (*D. fimbriatus*)

We used the data from the four *D. fimbriatus* populations to assess the effect of latitude, and called the model “modClim” in the following. We modelled the SCP with several candidate linear models including predictor variables Diff (time between capture and SCP measurements), site, climate (continental/temperate, as defined by the biogeographic location), sex and body size. We also considered the interaction between climate and body size and/or the interaction between body size and site (See appendix 1 for the list of candidate models).

### Comparison of SCP between species (northern populations)

We used *D. fimbriatus* and *D. plantarius* from Scandinavia to compare the SCP of species from northern populations, and called the model modSp in the following. We modelled the SCP with several candidate linear models with variables Diff, site, species, sex and body size, as well as the interaction between species and body size and/or the interaction between body size and site (See appendix 2 for the list of candidate models).

### Statistical Analysis

We used packages rstanarm (Goodrich et al. 2020), modelbased (Makowski et al. 2020) and bayestestR (Makowski et al. 2019a) in R (R Core Team 2020) to fit the linear models in a Bayesian framework. We used a normal distribution centred on 0 and a standard deviation of 2.5 as weakly informative priors (rather than using flat priors, see Gelman et al. 2008, Gelman and Shalizi 2013). We fitted the models using four chains and 4000 iterations. We used leave-one-out-cross-validation value (LOO value) to compare the predictive accuracy of fitted models, and to select the most accurate model (Vehtari et al. 2017). We checked the convergence of the models both visually and by making sure that Rhat value was not larger than 1.1 (Gelman and Rubin 1992).

Following Makowski et al. (2019b), we used the probability of direction (pd), which is the probability that the posterior distribution of a parameter is strictly positive or negative, to describe the existence of an effect of an explanatory variable. We used the percentage of the full region of practical equivalence (ROPE) lower than 1% as an index of the significance of an effect. We represented the uncertainty with a credible interval of 89%.

## Results

### General results

The SCP of the spiders varied from −2.6 to −16.4°C, with an average of −7.8±2.3°C (n=119). Fig. 2 shows typical cooling curves of *Dolomedes fimbriatus* (from C2) and *Dolomedes plantarius* (from C3) with exotherms of about 8 and 6.5°C and a SCP of −9.3 and −7.5°C respectively. None of the spiders tested survived freezing.

The body size of juveniles of *D. plantarius* was on average 5.36±0.69mm while body size of *D. plantarius* was 4.28±0.56mm in the South and 4.53±0.47mm in the South.

### Validation and selection of models

All of our candidate models converged (Rhat < 1.1). According to LOO values, some models were considered equivalent (Appendices 1 & 2). The modClim model with the lowest LOO value and therefore the highest predictive power included variables Diff (time between capture and test), climate, body size and the interactive effect of climate and body size (table 2). For modSp model, the best model included Diff, species, body size and the interactive effect of body size and species.

**Table 2:** Parameter estimates of the most accurate model explaining the SCP values between different climatic areas for D. fimbriatus (modClim, see appendix 1). CI: 89% credible intervals, pd: probability of direction, ROPE: percentage of the full region of practical equivalence. Diff: time difference between date of capture and date of test; Temperature: climate variable (continental climate in the intercept); Temperate:Body size: interactive effect of the climate and body size.

|  | Estimate | CI low | CI high | pd | ROPE (%) | Rhat |
| --- | --- | --- | --- | --- | --- | --- |
| (Intercept) | -8.3 | -12.3 | -4.7 | 1.00 | 0.0 | 1.0 |
| Diff | -0.3 | -0.4 | -0.2 | 1.00 | 23.1 | 1.0 |
| Temperate | 8.1 | 3.2 | 12.8 | 0.99 | 0.4 | 1.0 |
| Body size | 8.4 | 0.6 | 16.5 | 0.96 | 0.9 | 1.0 |
| Temperate:Body size | -14.3 | -24.7 | -2.8 | 0.98 | 0.4 | 1.0 |

### Comparison of SCP across latitudes (*D. fimbriatus*)

Regarding modClim (table 2), the SCP of individuals of southern and northern populations significantly differed (pd = 99%, <1% in ROPE, fig. 3) and were −6.6±2.3°C (min. −11.5°C, max. −2.6°C; n=50) and −9.05±2.31°C (min. −6.30°C, max. −2.30°C, n=48), respectively. The SCP significantly increased with the spider’s body size (pd = 96%, <1% in ROPE, median = 8.4 [0.6; 16.5]), which means that larger spiders had higher SCP than smaller spiders. The effect of spiders’ body size on the SCP was significantly different between the two climatic areas (pd = 98%, <1% in ROPE; fig. 4). Namely, the SCP increased with the body size of spiders in the northern climate (pd = 96%, <1% in ROPE, median = 8.33 [0.15; 15.56]) while the relation between SCP and body size in the South was not different from 0 (<1% in ROPE but pd < 90%).

**Figure 3:**
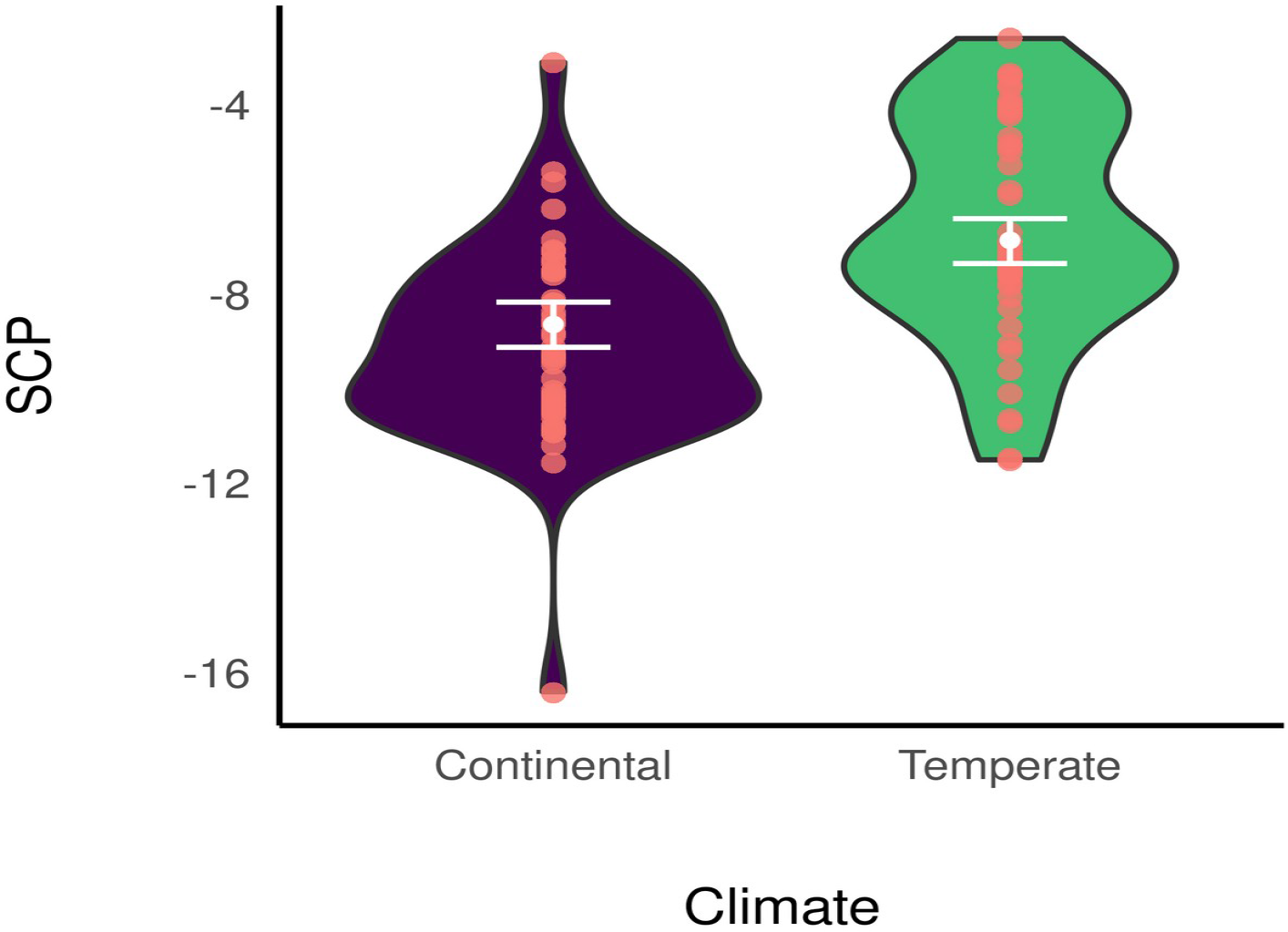
Marginal posterior means of SCP (white dot) estimated under modClim for the two different climatic areas and its 89% credible interval (white bar). Red dots represent the original data and the violin distributions represent a density plot.

**Figure 4:**
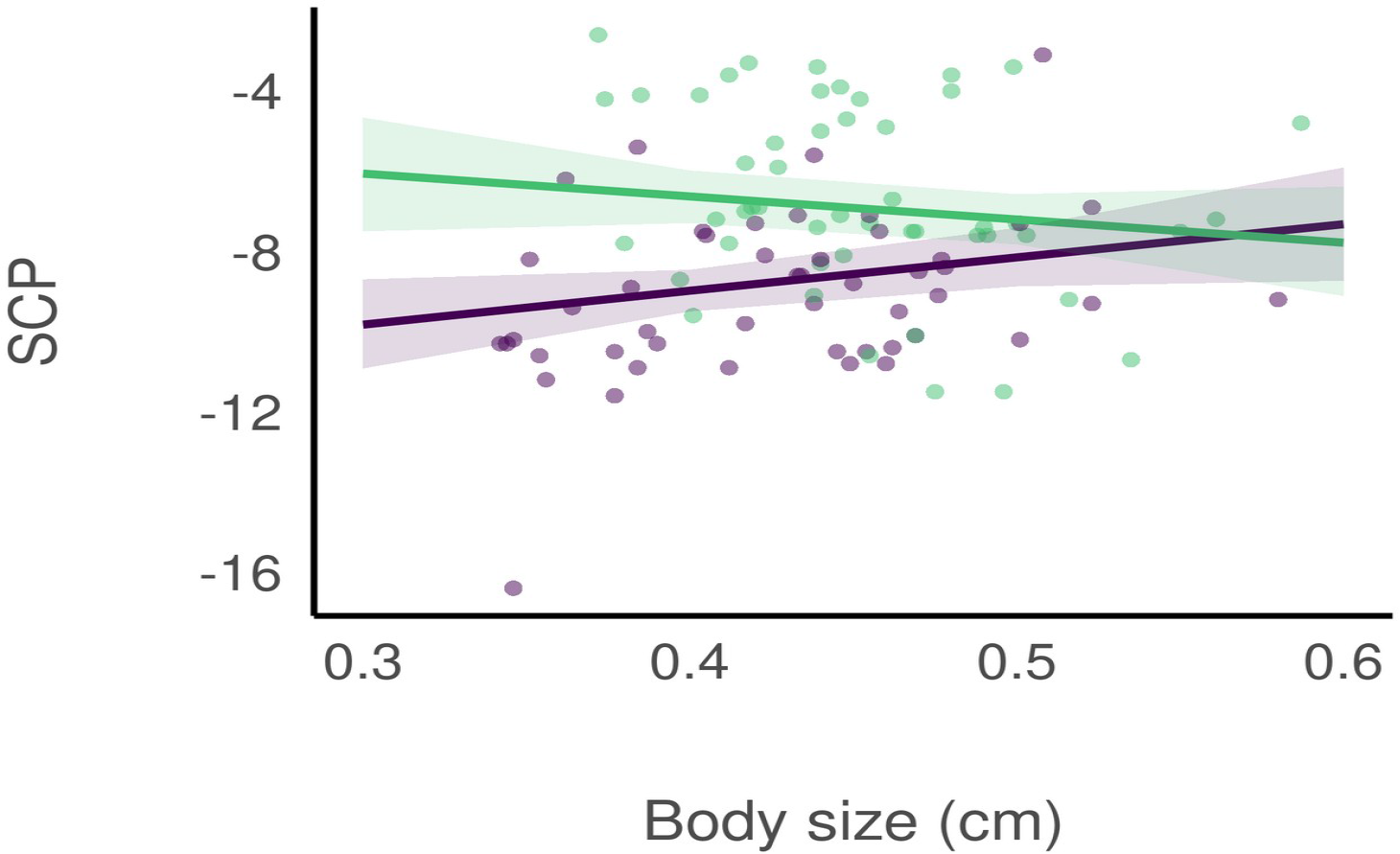
Predicted effect of *D. fimbriatus* body size on the SCP, and its 89% credible interval, for the two different climatic areas under modClim. Purple: predictions for the continental climate, green: predictions for the temperate climate; dots represent original data.

### Comparison of SCP between species (northern populations)

Regarding modSp (table 3), the SCP significantly increased with body size of both species together (pd = 98%, <1% in ROPE, Median = 9.1 [2.0; 15.9]; fig. 5). Nonetheless, the effect of body size on the SCP was not different between species (pd = 93%). The SCPs of individuals of *D. plantarius* and *D. fimbriatus* of northern populations likely differed (pd = 95%, 1%<ROPE<2.5%) and was −7.56±0.32 (min. −9.4°C, max. −4.4°C; n=21) for *D. plantarius* (for *D. fimbriatus*, see above). We did not find a significant effect of Diff for modSp (ROPE = 23%).

**Table 3:** Parameter estimates of the most accurate model explaining the SCP values between the two species in continental climate (modSp, see appendix 2). CI: 89% credible intervals, pd: probability of direction, ROPE: percentage of the full region of practical equivalence. Diff: time difference between date of capture and date of test; *D. plantarius*: species variable (D. fimbriatus in the intercept); *D. plantarius*:Body size: interactive effect of species and body size.

|  | Estimate | CI low | CI high | pd | ROPE (%) | Rhat |
| --- | --- | --- | --- | --- | --- | --- |
| (Intercept) | -9.77 | -13.8 | -5.6 | 1.00 | 0.0 | 1.0 |
| Diff | -0.22 | -0.4 | -0.0 | 0.97 | 42.8 | 1.0 |
| *D. plantarius* | 4.80 | 0.1 | 9.4 | 0.95 | 1.6 | 1.0 |
| Body size | 9.10 | 2.0 | 15.9 | 0.98 | 0.5 | 1.0 |
| *D. plantarius*:Body size | -8.65 | -17.4 | 1.0 | 0.93 | 0.8 | 1.0 |

**Figure 5:**
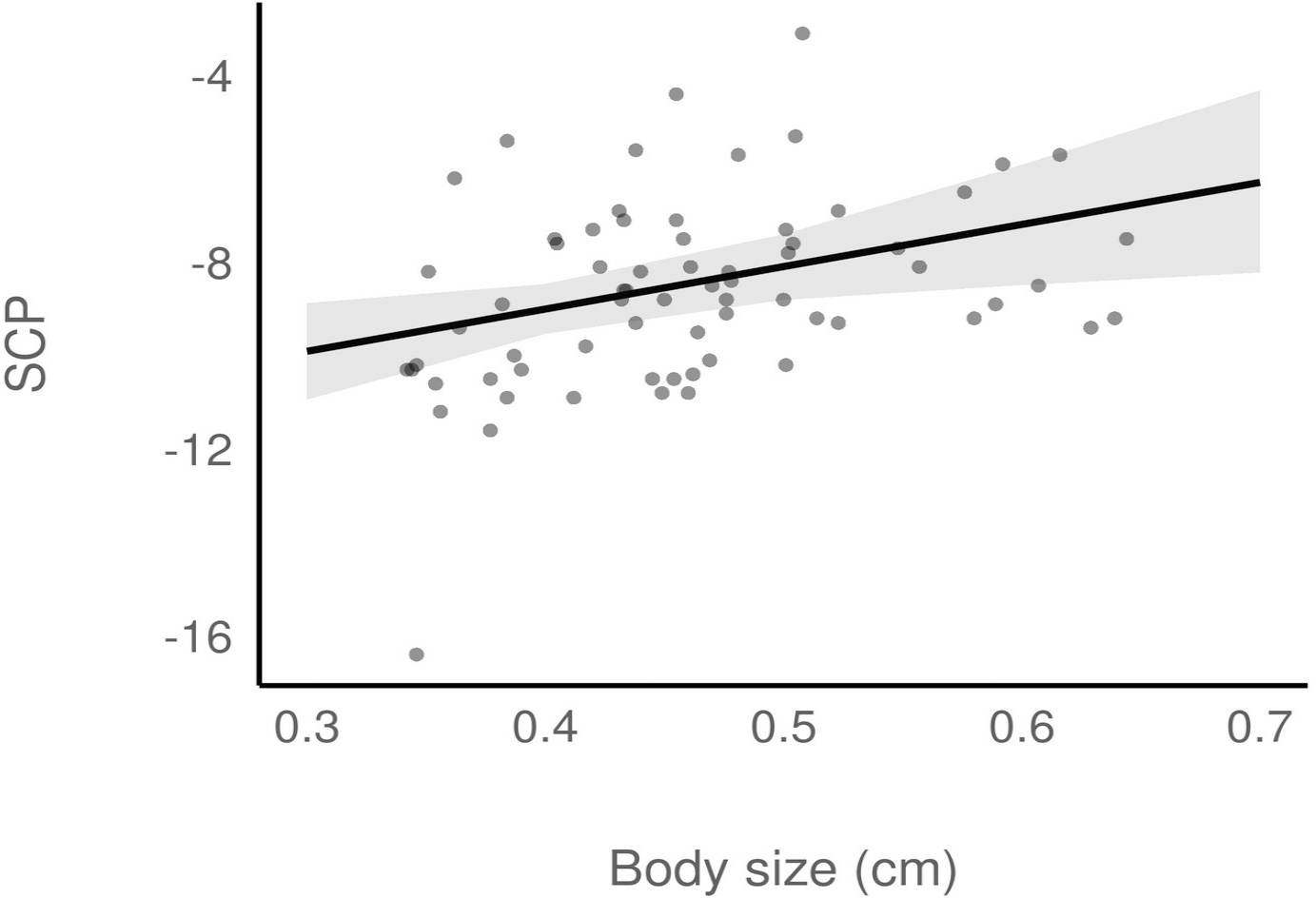
Predicted effect of body size of *D. plantarius* and *D. fimbriatus* on the SCP in Scandinavia, and its 89% credible interval, under modSp. Dots represent original data.

## Discussion

Our study showed that the SCP of northern fishing spiders from a continental climate was lower than the SCP of southern *Dolomedes* from a temperate climate. The SCP was positively related to body size for both species, but this relationship differed between the two climates for *D. fimbriatus*. Finally, we found that the SCP of *Dolomedes fimbriatus* was slightly lower than that of *Dolomedes plantarius*.

The SCP of *D. fimbriatus* decreased with increasing latitude, while juveniles of the species did not differ in size. In this study, we tested four populations from two biogeographic locations which characterised different climates and latitudes along the species distribution range. The northern populations, at the range limit, experience cold winters with permanent snow cover, whereas the southern populations, from a more central latitude of the range, experience warmer winters with only rarely a snow cover. The northern and southern locations are characterised by temperate and continental climate respectively (Kottek et al. 2006) and the corresponding range of temperatures might explain the decrease in SCP towards the North. Indeed, temperature influences cold hardiness in arthropods, including spiders (Nentwig 2012) and a poleward increase in thermal tolerance is observed in many ectotherms (Sunday et al. 2011). An acclimation to warmer temperatures, as for southern spiders, can also reduce the tolerance to cold conditions (Jensen et al. 2019). At the same time, northern spiders could benefit from their cold acclimation by being more active during cooler periods in summer (Everatt et al. 2013). The diurnal range also differs along the latitudinal gradient; i.e. northern populations stand more substantial variation in the diurnal activity range.

The impact of diurnal activity range, together with temperature, are essential cues to determine the cold resistance of ectotherm arthropods (e.g. soil dwelling collembolan *Orchesella cincta* see Jensen et al. 2019, or Paaijmans et al. 2013, Seebacher et al. 2015). These might have impacted spiders differently at the time of our experiments (late summer/ early autumn), as northern *Dolomedes* are confronted to earlier and harsher winter. These two cues have been shown to impact the overwintering of another *Dolomedes* species, from North America (*D. triton*; Spence and Zimmermann 1998), and might similarly impact the overwintering of *D. fimbriatus*. To our knowledge, *Dolomedes* species are inactive during winter (Aitchison 1984). Schmidt (1957) noted that *D. fimbriatus* overwinters twice before reaching the adult stage. He also noted that juveniles spend the winter in dry vegetation at high strata, which is probably the overwintering habitat of the southern spiders we tested here. However, the northern *Dolomedes* we tested endure temperatures colder than the SCP measured in this study. For this reason, we hypothesised that, similarly to *Dolomedes triton* in Canada (Spence and Zimmermann 1998), spiderlings and juveniles overwinter under the snow. Indeed, the temperature in the subnivean layer, which is between the soil surface and the base of the snowpack, is warmer and more stable than the air temperature above the snow, and protect species from temperatures lower than their SCP (Marchand 1982).

*Dolomedes*, like other spider species, are not freezing tolerant as none of the spiders tested survived freezing. Cold-hardiness of *Dolomedes* is important for winter survival. Based on the cold hardiness classification of Bale (1996) and Bale (2002) (see also appendix 3 for a summarised classification), we hypothesise that *Dolomedes*, at least from the northern populations, are either chill-susceptible or freeze-avoidant. The main difference between these two cold hardiness classes is the ability to survive damages caused by cold injuries. Freezing-avoidant species survive until freezing, while chill-tolerant die sooner due to chill injuries. Spiders from a close family (*Pardosa*, Lycosidae) at northern latitudes are from the same cold hardiness class (Anthony and Sinclair 2019). Nonetheless, we only tested the SCP and more measurements, such as the lower lethal temperature, would be necessary to define the cold hardiness class more precisely. The cold hardiness class of *Dolomedes* might also vary between the two biogeographic positions as demonstrated for the butterfly *Piries rapae* which is either freeze-tolerant or freezing-avoidant depending on the latitude (Li and Zachariassen 2007).

Even if *Dolomedes* from the two areas did not differ in body size, we found an overall decrease of the SCP with increasing spider body size. Smaller individuals being more cold tolerant than bigger ones is a general trend for ectotherm animals (e.g. for ants see Hahn et al. (2008), for beetles see Johnston and Lee (1990)). This trend is also observed for spiders with smaller instars being more tolerant to cold than larger juveniles and adults (Almquist 1970, Bayram and Luff 1993), and it might explain our extreme SCP measures down to −16.8 °C for one *D. fimbriatus* from Fennoscandia.

The decline in supercooling abilities with increasing latitude was nonetheless not observed in southern *D. fimbriatus*. The size of southern spiders, that have higher SCP, seemed to be less related to SCP. This difference in strategy between temperate and colder habitats has been reported in other species from the closely-related family of Lycosidae (Ameline et al. 2018). The northern spiders have a shortened breeding season, which can impact life history traits such as body size (Bowden et al. 2015). The smaller fishing spiders under continental climate could be advantaged as they can survive colder winters. After the winter, northern fishing spiders could accelerate their development because cold-adapted ectotherms have a higher metabolic rate in an environment with limited energy (Sinclair et al. 2012).

We found slightly higher resistance to cold temperature in *D. fimbriatus* compared to *D. plantarius* (for populations at similar latitudes), which might be partly due to the smaller size of *D. fimbriatus*. In turn, this difference between species might explain the wider northward distribution of *D. fimbriatus* compared to that of *D. plantarius*. It indeed appears that specialist species are larger under harsher conditions because they are more adapted to their environment (Ameline et al. 2018). A larger size implies a smaller cold resistance here, which might be detrimental in this case. Nonetheless, the SCPs measured in this study were close to those measured for phylogenetically close spiders (from the same Lycosoidea superfamily) from northern latitudes (Anthony and Sinclair 2019). These values are considered as medium cold resistance (Nentwig 2012).

Climate change impacts spiders in various ways. At northern latitudes, subnivean layer is supposedly a non-freezing environment with quite stable temperatures (Pruitt 1957) but snow density and length of the snow season impacts the stability of these conditions (Pauli et al. 2013, Bale and Hayward 2010). While air temperature increases with climate change, the subnivean layer may become colder (Wipf and Rixen 2010). This paradox is already negatively impacting invertebrates (Williams et al. 2015, Slatyer et al. 2017). Even though we found that fishing spiders from continental climate tolerated colder temperatures than spiders from temperate climate, the lowest SCP was higher than the lowest air temperature measured historically in Fennoscandia. A weakened subnivean shelter could negatively impact northern populations and even more so for the rare *D. plantarius* which is less cold resistant. Another impact of the increased length of the snow free season could be a second clutch in northern *Dolomedes*, as reported in the arctic Lycosidae *Pardosa glacialis* (Høye et al. 2020).

We found that the cold tolerance of fishing spiders varied among populations, between climates and between species. Nonetheless, the difference in SCP between the two species was not striking. Sample another population of *D. plantarius* could support the slight difference found between species, but we tried to limit the impact of sampling on populations of this red-listed species (World Conservation Monitoring Centre 1996). Moreover, we assessed cold tolerance based on measuring the SCP only and from spiders sampled in late summer / early autumn. Sampling *Dolomedes* is challenging especially at northern latitudes in winter. Nonetheless, studying life history traits like cold resistance is valuable to explore and predict the distribution of understudied invertebrates (Mammola et al. 2020), especially by integrating ecophysiology of species.

## References

Addo-Bediako, A., S. L. Chown, and K. J. Gaston. 2000. Thermal tolerance, climatic variability and latitude. Proceedings of the Royal Society B: Biological Sciences 267:739–745.

Aitchison, C. W. 1984. Low Temperature Feeding by Winter-Active Spiders. Journal of Arachnology 12:297–305.

Almquist, S. 1970. Thermal tolerances and preferences of some dune-living spiders. Oikos 21:230–235.

Ameline, C., C. Puzin, J. J. Bowden, K. Lambeets, P. Vernon, and J. Pétillon. 2018. Habitat specialization and climate affect arthropod fitness: A comparison of generalist vs. Specialist spider species in Arctic and temperate biomes. Biological Journal of the Linnean Society 121:592–599.

Ansart, A., A. Guiller, O. Moine, M.-C. Martin, and L. Madec. 2014. Is cold hardiness size-constrained? A comparative approach in land snails. Evolutionary Ecology 28:471–493.

Anthony, S. E., C. M. Buddle, T. T. Høye, and B. J. Sinclair. 2019. Thermal limits of summer-collected *Pardosa* wolf spiders (Araneae: Lycosidae) from the Yukon Territory (Canada) and Greenland. Polar Biology 42:2055–2064.

Anthony, S. E., and B. J. Sinclair. 2019. Overwintering red velvet mites are freeze tolerant. Physiological and Biochemical Zoology 92:201–205.

Bale, J. S. 1996. Insect cold hardiness: A matter of life and death. European Journal of Entomology 93:369–382.

Bale, J. S. 2002. Insects and low temperatures: From molecular biology to distributions and abundance. Philosophical Transactions of the Royal Society of London. Series B: Biological Sciences 357:849–862.

Bale, J. S., and S. a. L. Hayward. 2010. Insect overwintering in a changing climate. Journal of Experimental Biology 213:980–994.

Bayram, A., and M. L. Luff. 1993. Cold-hardiness of wolf-spiders (Lycosidae, Araneae) with particular reference to *Pardosa pullata* (Clerck). Journal of thermal biology.

Blanckenhorn, W. U., and M. Demont. 2004. Bergmann and converse Bergmann latitudinal clines in arthropods: Two ends of a continuum? Integrative and Comparative Biology 44:413–424.

Bowden, J. J., R. R. Hansen, K. Olsen, and T.T. Høye. 2015. Habitat-specific effects of climate change on a low-mobility Arctic spider species. Polar Biology 38:559– 568.

Colinet, H., P. Vernon, and T. Hance. 2007. Does thermal-related plasticity in size and fat reserves influence supercooling abilities and cold-tolerance in *Aphidius colemani* (Hymenoptera: Aphidiinae) mummies? Journal of Thermal Biology 32:374–382.

Danks, H. V. 2004. Seasonal adaptations in Arctic insects. Integrative and Comparative Biology 44:85–94.

David, J.-F., M.-L. Célérier, and G. Vannier. 1996. Overwintering with a low level of cold-hardiness in the temperate millipede *Polydesmus angustus*. Acta Oecologica 17:393–404.

David, J.-F., and G. Vannier. 1996. Changes in supercooling with body size, sex, and season in the long-lived milliped *Polyzonium germanicum* (Diplopoda, Polyzonidae). Journal of Zoology 240:599–608.

Ditrich, T., V. Janda, H. Vaněčková, and D. Doležel. 2018. Climatic variation of supercooling point in the Linden Bug *Pyrrhocoris apterus* (Heteroptera: Pyrrhocoridae). Insects 9:144.

Duman, J. G. 2001. Antifreeze and ice nucleator proteins in terrestrial arthropods. Annual Review of Physiology 63:327–357.

Duman, J. G., V. Bennett, T. Sformo, R. Hochstrasser, and B. M. Barnes. 2004. Antifreeze proteins in Alaskan insects and spiders. Journal of Insect Physiology 50:259–266.

Everatt, M. J., J. S. Bale, P. Convey, M. R. Worland, and S. A. L. Hayward. 2013. The effect of acclimation temperature on thermal activity thresholds in polar terrestrial invertebrates. Journal of Insect Physiology 59:1057–1064.

Fick, S. E., and R. J. Hijmans. 2017. WorldClim 2: New 1-km spatial resolution climate surfaces for global land areas. International Journal of Climatology 37:4302– 4315.

Gaston, K. J. 2003. The structure and dynamics of geographic ranges. Oxford University Press.

Gelman, A., A. Jakulin, M. G. Pittau, and Y.-S. Su. 2008. A weakly informative default prior distribution for logistic and other regression models. Annals of Applied Statistics 2:1360–1383.

Gelman, A., and D. B. Rubin. 1992. Inference from iterative simulation using multiple sequences. Statistical Science 7:457–472.

Gelman, A., and C. R. Shalizi. 2013. Philosophy and the practice of Bayesian statistics. British Journal of Mathematical and Statistical Psychology 66:8–38.

Goodrich, B., J. Gabry, I. Ali, and S. Brilleman. 2020. Rstanarm: Bayesian applied regression modeling via Stan.

Guarisco, H. 2010. The Fishing Spider genus *Dolomedes* (Araneae: Pisauridae) in Kansas. Transactions of the Kansas Academy of Science 113:35–43.

Hahn, D. A., A. R. Martin, and S. D. Porter. 2008. Body Size, but Not Cooling Rate, Affects Supercooling Points in the Red Imported Fire Ant, *Solenopsis invicta*. Environmental Entomology 37:1074–1080.

Høye, T. T., J.-C. Kresse, A. M. Koltz, and J. J. Bowden. 2020. Earlier springs enable high-Arctic wolf spiders to produce a second clutch. Proceedings of the Royal Society B: Biological Sciences 287:20200982.

Jakob, E. M., S. D. Marshall, and G. W. Uetz. 1996. Estimating fitness: A comparison of body condition indices. Oikos 77:61–67.

Jensen, A., T. Alemu, T. Alemneh, C. Pertoldi, and S. Bahrndorff. 2019. Thermal acclimation and adaptation across populations in a broadly distributed soil arthropod. Functional Ecology 33:833–845.

Johnston, S. L., and R. E. Lee. 1990. Regulation of supercooling and nucleation in a freeze intolerant beetle (*Tenebrio Molitor*). Cryobiology 27:562–568.

Kirchner, W. 1973. Ecological aspects of cold resistance in spiders (a comparative study). Pages 271–279 in . Wieser, editor. Effects of temperature on ectothermic organisms: Ecological implications and mechanisms of compensation. Springer, Berlin, Heidelberg.

Kottek, M., J. Grieser, C. Beck, B. Rudolf, and F. Rubel. 2006. World Map of the Köppen-Geiger climate classification updated. Meteorologische Zeitschrift 15:259–263.

Lee, R. E., and J. P. Costanzo. 1998. Biological ice nucleation and ice distribution in cold-hardy ectothermic animals. Annual Review of Physiology 60:55–72.

Li, N., and K. Zachariassen. 2007. Cold hardiness of insects distributed in the area of Siberian Cold Pole. Comparative Biochemistry and Physiology Part A: Molecular & Integrative Physiology 146:S156–S157.

Makowski, D., M. Ben-Shachar, and D. Lüdecke. 2019a. bayestestR: Describing effects and their uncertainty, existence and significance within the Bayesian framework. Journal of Open Source Software 4:1541.

Makowski, D., M. S. Ben-Shachar, S. H. A. Chen, and D. Lüdecke. 2019b. Indices of effect existence and significance in the Bayesian framework. Frontiers in Psychology 10.

Makowski, D., D. Lüdecke, and M. S. Ben-Shachar. 2020. Modelbased: Estimation of model-based predictions, contrasts and means. CRAN.

Mammola, S., J. Pétillon, A. Hacala, J. Monsimet, S.-l. Marti, P. Cardoso, and D. Lafage. 2020. Challenges and opportunities of species distribution modelling of terrestrial arthropods. [Manuscript submitted for publication].

Marchand, P. J. 1982. An index for evaluating the temperature stability of a subnivean environment. The Journal of Wildlife Management 46:518–520.

Monsimet, J., O. Devineau, J. Pétillon, and D. Lafage. 2020. Explicit integration of dispersal-related metrics improves predictions of SDM in predatory arthropods. Scientific Reports 10:1–12.

Mousseau, T. A. 1997. Ectotherms follow the converse to Bergmann’s rule. Evolution 51:630–632.

Murphy, J., T. Rossolimo, and S. Adl. 2008. Cold-hardiness in the wolf spider *Pardosa groenlandica* (Thorell) with respect to thermal limits and dehydration. The Journal of Arachnology 36:213–215.

Nentwig, W. 2012. Ecophysiology of spiders. Springer Science & Business Media.

Overgaard, J., and H. A. MacMillan. 2017. The integrative physiology of insect chill tolerance. Annual Review of Physiology 79:187–208.

Paaijmans, K. P., R. L. Heinig, R. A. Seliga, J. I. Blanford, S. Blanford, C. C. Murdock, and M. B. Thomas. 2013. Temperature variation makes ectotherms more sensitive to climate change. Global Change Biology 19:2373–2380.

Pauli, J. N., B. Zuckerberg, J. P. Whiteman, and W. Porter. 2013. The subnivium: A deteriorating seasonal refugium. Frontiers in Ecology and the Environment 11:260–267.

Penell, A., F. Raub, and H. Höfer. 2018. Estimating biomass from body size of European spiders based on regression models. Journal of Arachnology 46:413– 419.

Pétillon, J., W. Montaigne, and D. Renault. 2009. Hypoxic coma as a strategy to survive inundation in a salt-marsh inhabiting spider. Biology Letters 5:442–445.

Pruitt, W. O. J. 1957. Observations on the bioclimate of some taiga mammals. Artic 10:130–138.

R Core Team. 2020. R: A Language and Environment for Statistical Computing. R Foundation for Statistical Computing, Vienna, Austria.

Renault, D., C. Salin, G. Vannier, and P. Vernon. 2002. Survival at low temperatures in insects: What is the ecological significance of the supercooling point? CryoLetters 23:217–228.

Schmidt, G. 1957. Einige notizen über *Dolomedes fimbriatus* (Cl.). Zoologischer Anzeiger 158:83–97.

Schneider, C. A., W. S. Rasband, and K. W. Eliceiri. 2012. NIH Image to ImageJ: 25 years of image analysis. Nature Methods 9:671–675.

Schulte, P. M., T. M. Healy, and N. A. Fangue. 2011. Thermal performance curves, phenotypic plasticity, and the time scales of temperature exposure. Integrative and Comparative Biology 51:691–702.

Seebacher, F., C. R. White, and C. E. Franklin. 2015. Physiological plasticity increases resilience of ectothermic animals to climate change. Nature Climate Change 5:61–66.

Sinclair, B. J., A. Addo-Bediako, and S. L. Chown. 2003. Climatic variability and the evolution of insect freeze tolerance. Biological Reviews 78:181–195.

Sinclair, B. J., L. E. C. Alvarado, and L. V. Ferguson. 2015. An invitation to measure insect cold tolerance: Methods, approaches, and workflow. Journal of Thermal Biology 53:180–197.

Sinclair, B. J., A. G. Gibbs, W.-K. Lee, A. Rajamohan, S. P. Roberts, and J. J. Socha. 2009. Synchrotron x-ray visualisation of ice formation in insects during lethal and non-lethal freezing. PLOS ONE 4:1–10.

Sinclair, B. J., C. M. Williams, and J. S. Terblanche. 2012. Variation in thermal performance among insect populations. Physiological and biochemical zoology 85:594–606.

Slatyer, R. A., M. A. Nash, and A. A. Hoffmann. 2017. Measuring the effects of reduced snow cover on Australia’s alpine arthropods. Austral Ecology 42:844– 857.

Smith, H. 2000. The Status and Conservation of the Fen Raft Spider (*Dolomedes plantarius*) at Redgrave and Lopham Fen National Nature Reserve, England. Biological Conservation 95:153–164.

Somero, G. N. 2012. The physiology of global change: Linking patterns to mechanisms. Annual Review of Marine Science 4:39–61.

Spence, J. R., and M. Zimmermann. 1998. Phenology and life-cycle of the fishing spider *Dolomedes Triton* Walckenaer (Araneae, Pisauridae) in Central Alberta. Canadian Journal of Zoology 76:295–309.

Spicer, J., and K. J. Gaston. 1999. Physiological diversity and Its ecological implications. Blackwell Science. Oxford Academic, Oxford.

Sunday, J. M., A. E. Bates, and N. K. Dulvy. 2011. Global analysis of thermal tolerance and latitude in ectotherms. Proceedings of the Royal Society B: Biological Sciences 278:1823–1830.

Sømme, L. 1982. Supercooling and winter survival in terrestrial arthropods. Comparative Biochemistry and Physiology Part A: Physiology 73:519–543.

Vehtari, A., A. Gelman, and J. Gabry. 2017. Practical Bayesian model evaluation using leave-one-out cross-validation and WAIC. Statistics and Computing 27:1413– 1432.

Vernon, P., and G. Vannier. 2002. Evolution of freezing susceptibility and freezing tolerance in terrestrial arthropods. Comptes Rendus Biologies 325:1185–1190.

Voorhies, W. A. V. 1996. Bergmann size clines: A simple explanation for their occurrence in ectotherms. Evolution 50:1259–1264.

Williams, C. M., H. A. L. Henry, and B. J. Sinclair. 2015. Cold truths: How winter drives responses of terrestrial organisms to climate change. Biological Reviews 90:214–235.

Wipf, S., and C. Rixen. 2010. A review of snow manipulation experiments in Arctic and alpine tundra ecosystems. Polar Research 29:95–109.

World Conservation Monitoring Centre. 1996. The IUCN Red List of Threatened Species 1996.

