## Appendix 1 for "Biogeographic position and body size jointly set lower thermal limits of wandering spiders"

Appendix 1: Comparison of the expected predictive accuracy of models comparing the two geographic areas (modClim). Difference of expected log predictive density and standard error between model for the leave-one-out cross validation (LOO) are shown (epld: expected log predictive density, SE: Standard-error of the epld).

| Code | Model | elpd_diff | se_diff |
|---|---|---|---|
| mC11 | SCP ~ Diff + Climate + Length + Climate:Length | 0.00 | 0.00 |
| mC8 | SCP ~ Diff + Climate + Sex + Length + Climate:Length | -0.10 | 1.20 |
| mC3 | SCP ~ Diff + Climate | -1.55 | 1.91 |
| mC10 | SCP ~ Diff + Site + Climate + Sex + Length + Climate:Length + Site:Length | -2.91 | 2.18 |
| mC4 | SCP ~ Diff + Site + Climate | -2.99 | 3.04 |
| mC2 | SCP ~ Diff + Site | -3.00 | 2.94 |
| mC5 | SCP ~ Diff + Site + Climate + Sex | -3.13 | 3.15 |
| mC12 | SCP ~ Diff + Site + Length + Site:Length | -3.15 | 1.83 |
| mC9 | SCP ~ Diff + Site + Sex + Length + Site:Length | -3.46 | 2.28 |
| mC6 | SCP ~ Diff + Site + Climate + Length | -3.87 | 2.74 |
| mC7 | SCP ~ Diff + Site + Climate + Sex + Length | -3.99 | 2.93 |
| mC1 | SCP ~ 1+Diff | -21.01 | 6.52 |
