## Appendix 2 for "Biogeographic position and body size jointly set lower thermal limits of wandering spiders"

Appendix 2: Comparison of the expected predictive accuracy of models comparing the two species (modSp). Difference of expected log predictive density and standard error between models for the leave-one-out cross validation (LOO) are shown (ELPD: expected log predictive density, SE: Standard-error of the ELPD).

| Code | Model | elpd_diff | se_diff |
|---|---|---|---|
| mS11 | SCP ~ Diff + Species + Length + Species:Length | 0.00 | 0.00 |
| mS8 | SCP ~ Diff + Species + Sex + Length + Species:Length | -0.02 | 1.33 |
| mS3 | SCP ~ Diff + Species | -1.46 | 2.31 |
| mS12 | SCP ~ Diff + Site + Length + Site:Length | -1.67 | 0.86 |
| mS10 | SCP ~ Diff + Site + Species + Sex + Length + Species:Length + Site:Length | -2.10 | 1.65 |
| mS9 | SCP ~ Diff + Site + Sex + Length + Site:Length | -2.23 | 1.57 |
| mS5 | SCP ~ Diff + Site + Species + Sex | -2.40 | 2.61 |
| mS6 | SCP ~ Diff + Site + Species + Length | -2.41 | 1.43 |
| mS7 | SCP ~ Diff + Site + Species + Sex + Length | -2.43 | 1.85 |
| mS2 | SCP ~ Diff + Site | -2.51 | 2.33 |
| mS4 | SCP ~ Diff + Site + Species | -2.72 | 2.40 |
| mS1 | SCP ~ 1+Diff | -5.33 | 4.23 |
