## Appendix 3 for "Biogeographic position and body size jointly set lower thermal limits of wandering spiders"

Appendix 3: Cold hardiness classes of exotherm arthropods. LLT: lower lethal temperature, SCP = supercooling point (Bale 1996 and 2002).

| Class | Subclass | Definition |
|---|---|---|
| Freezing tolerant | | LLT < SCP |
| Freezing avoidant | Freeze-avoidant | LLT = SCP |
| | Chill-tolerant | LLT ≥ SCP |
| | Chill-susceptible | LLT > SCP |
| | Opportunistic-survival | LLT ≫ SCP |
